# Biochemical characterization of ABHD14A, an outlying member of the metabolic serine hydrolase family

**DOI:** 10.1101/2025.11.28.691245

**Authors:** Sonali Gupta, Siddhesh S. Kamat

## Abstract

Certain uncharacterised members of the metabolic serine hydrolase enzyme family remain difficult to annotate due to poor tractability, context-dependent expression, and the absence of defined biochemical activities. Here, we provide the first functional characterization of the human enzyme ABHD14A. By engineering a soluble N-terminally truncated variant, we demonstrate by gel-based activity-based protein profiling and p-nitrophenyl-ester hydrolysis assays that ABHD14A is an active enzyme that preferentially turns over short-chain esters. Notably, ABHD14A exhibits a CoA-dependent enhancement of p-nitrophenyl-acetate hydrolysis, indicative of a ping-pong type acetyltransferase mechanism similar to that previously described for another homologous ABHD14-enzyme, ABHD14B. To investigate endogenous protein levels, we generated a high-affinity polyclonal antibody and found that ABHD14A is undetectable across a panel of immortalized mammalian cell lines and adult mouse tissues, challenging current transcriptomic database predictions. Upon heterologous expression in HEK293T cells, full length ABHD14A is catalytically active and localizes specifically to the Golgi apparatus, suggesting a specialized role in secretory pathway biology. Together, these findings establish the enzymatic activity, mechanistic features, and subcellular localization of ABHD14A, while providing essential biochemical and immunological tools that now enable the systematic discovery of its physiological substrates and regulatory contexts.

## INTRODUCTION

Uncharacterised human proteins remain a significant frontier in contemporary molecular and cellular biology, with a substantial fraction of the human proteome still lacking definitive functional annotation^1–3^. This gap persists despite major advances in genomics, proteomics, and machine learning–driven prediction, reflecting the inherent difficulty of assigning biochemical roles to proteins whose expression patterns are often transient, cell-type specific, or activated only under particular physiological states. Increasing evidence underscores that protein function cannot be fully understood from sequence alone^4^; instead, many proteins are subject to tightly regulated, context-dependent expression influenced by developmental stage, metabolic status, immune activation, or environmental stress. Such dynamic regulation can obscure the detection of protein activities and hinder targeted studies, contributing to the slow pace of characterisation across large enzyme families and metabolic networks.

These challenges are quite evident within the metabolic serine hydrolase (mSH) superfamily – one of the largest, most functionally diverse, and still incompletely annotated enzyme groups in humans^5^. The mSHs mediate essential biochemical processes, including lipid metabolism, ester and amide hydrolysis, intercellular signalling, neurotransmission, and detoxification^5^. Yet several members of this superfamily lack well-defined substrates and physiological roles. Functional redundancy, overlapping substrate preferences, and subtle phenotypes upon genetic perturbation complicate efforts to infer physiological relevance. While the chemical proteomics technique, activity-based protein profiling (ABPP), has significantly enabled mapping enzymatic functions within the mSH family across tissues and disease states, activities for several enzymes from this enzyme family still remain elusive^6–8^. As a result, the complete annotation of this enzyme family remains incomplete, highlighting the need for more targeted studies that integrate structural, biochemical, and perhaps, even cellular evidence.

Within this context, the enzyme ABHD14A stands out as a markedly understudied protein from the mSH family whose biochemical activity remains poorly defined. Bioinformatics studies show that this enzyme contains a conserved catalytic serine within the canonical hydrolase motif [α/β-hydrolase domain (ABHD)], and structural models predict an active-site architecture consistent with ester or amide hydrolysis^9^. Yet, no definitive endogenous substrate, pathway, or physiological role has been assigned to this enzyme. Hence, as such, ABHD14A represents an interesting opportunity to elucidate how uncharacterised mSHs might integrate into human physiology. On this note, a deeper understanding of its expression control, substrate specificity, and structural determinants could illuminate not only the function of this single enzyme but also provide broader insights into generic strategies for annotating neglected proteins within large enzymatic families, such as the mSHs.

In this paper, we recombinantly purify a truncated variant of human ABHD14A from *E. coli*, and biochemically assay this enzyme. We demonstrate using gel-based ABPP and colorimetric assays, that human ABHD14A is indeed an active enzyme, capable of hydrolysing p-nitrophenyl-esters. We also show that human ABHD14A possesses an acetyltransferase activity with Coenzyme A (Co-A) being the other substrate, and acetyl-Co-A being the product of this biochemical reaction. To investigate the expression of ABHD14A in mammalian cells and tissues, we develop a polyclonal anti-ABHD14A antibody, and show contrary to publicly available databases, that most immortalized cell lines and adult mouse tissues lack ABHD14A expression. Finally, we overexpress full length ABHD14A in HEK293T cells, and show that this enzyme in active in mammalian cells and is sub-cellularly localized to the Golgi. Overall, our studies provide the first evidence of any activity for this cryptic enzyme, and open new avenues to study this poorly characterized enzyme in the context of mammalian physiology.

## MATERIALS AND METHODS

### Materials

All chemicals, buffers, and reagents were purchased from Sigma-Aldrich, and all tissue culture media and consumables were purchased from HiMedia, unless mentioned otherwise.

### Cloning and expression of recombinant human ABHD14A in *E. coli*

The wild type (WT) human *abhd14a* gene (UniProt ID: Q9BUJ0) was synthesized from GenScript as a codon-optimized construct for expression in *E. coli*. The human ABHD14A contains 271 amino acids, including an approximately 60-residue N-terminal transmembrane segment. Because full-length protein expression and purification was unsuccessful, a series of N-terminally truncated constructs (successive 10-aa deletions) bearing an N-terminal 6X-His tag were cloned into the pET45b(+) vector (Millipore). Recombinant plasmids were transformed into *E. coli* BL21(DE3). Cells were lysed in 1X-phosphate buffered saline (PBS), and soluble and membrane fractions were separated by centrifugation (21,000g, 4 °C, 45 min). Equal protein amounts from each fraction were analysed by SDS-PAGE to evaluate expression and solubility.

### Purification of ΔN60-ABHD14A variants

*E. coli* BL21(DE3) containing the N-terminal 6X-His tagged ΔN60-WT ABHD14A construct was grown in LB medium with 100 µg/mL ampicillin at 37 °C with constant shaking. At an optical density of 0.6 at 600 nm (OD_600_ ∼ 0.6), protein expression was induced by adding 100 µM isopropyl-β-*D*-1-thiogalactopyranoside and reducing the temperature to 18 °C for 16 h. Cells were harvested by centrifugation (6000 rpm, 20 min, 4 °C) and resuspended in lysis buffer (50 mM Tris, 50 mM imidazole, 500 mM NaCl at pH 8.0). Lysates were prepared by sonication, and clarified by centrifugation (15,000g, 4 °C, 45 min). The resulting soluble fraction was applied to a Ni²⁺-NTA affinity column (GE Healthcare), which was pre-equilibrated with the lysis buffer. The protein bound to the column was eluted with an elution buffer (50 mM Tris, 250 mM imidazole, 500 mM NaCl at pH 8.0) as per manufacturer’s instructions. Eluted fractions were dialyzed in the assay buffer (50 mM Tris, 500 mM NaCl at pH 8.0) to remove excess imidazole, flash-frozen in liquid nitrogen, and stored at −80 °C until further use. A catalytically inactive S171A variant of ABHD14A (N-terminal 6X-His tagged ΔN60-S171A ABHD14A) was generated using site directed mutagenesis (New England Biolabs) as per manufacturer’s instructions, and purified using the procedure mentioned above. Protein purity was assessed by SDS-PAGE analysis, and concentrations were determined using the Bradford assay.

### Gel based ABPP assays of the ΔN60-ABHD14A variants

All gel-based ABPP assays were done as per procedures previously reported by us^9, 10^, using the fluorophosphonate-rhodamine (FP-rhodamine) probe. For protein titrations, the ΔN60-ABHD14A variant (WT and S171A) concentrations ranged from 2 to 10 μM, with a constant FP-rhodamine concentration of 5 μM. For probe titration experiment, the ΔN60-ABHD14A variant (WT and S171A) concentration was maintained at 10 μM, and FP-rhodamine concentrations varied from 0.2 to 5 μM. Samples were resolved by 10% SDS-PAGE and the in-gel fluorescence of the probe-labelled proteins was visualized on the iBright1500 gel documentation system (Invitrogen). All gels were subsequently stained with Coomassie Brilliant Blue to confirm appropriate protein loading.

### p-Nitrophenyl-ester hydrolysis assays of the ΔN60-ABHD14A variants

All p-nitrophenyl-ester assays was performed using substrates and procedures previously reported by us^9, 10^, and the release of p-nitrophenol from the p-nitrophenyl-ester was monitored at 405 nm. A typical p-nitrophenyl-ester hydrolysis reaction was performed in a 250 μL volume, contained 10 μM ΔN60-WT ABHD14A [native or heat denatured (negative control)] and 500 μM p-nitrophenyl-ester in the assay buffer. To determine the effect of Co-A or Acetyl-Co-A on the rate of p-nitrophenyl-ester hydrolysis, ΔN60-WT ABHD14A (10 μM) was incubated with Co-A (1 mM) or Acetyl-Co-A (1 mM) at 37 °C for 15 min, and the reaction were initiated by adding 50 μM p-nitrophenyl-ester under assay conditions previously reported by us^10^.

### Production of an ABHD14A antibody

The anti-ABHD14A primary antibody was generated in rabbits using the purified human ΔN60-WT ABHD14A as the antigen in the National Facility for Gene Function in Health and Disease (NFGFHD) at IISER Pune using in house protocols [Formal approval from the IISER Pune – Institutional Animal Ethics Committee (IISER-P IAEC) (protocol no: IISER_Pune/ IAEC/2021_01/08 and IISER_Pune/ IAEC/2023_01/05)]. Briefly, a female New Zealand rabbit was immunized with 0.3 mg of purified human ΔN60-WT ABHD14A emulsified in Complete Freund’s Adjuvant (day 0). Thereafter, post-immunization, booster doses were administered on days 14, 42, 70 and 98. The serum was collected 14 days after each booster dose and terminally on day 126. The antibody titres from the harvested sera were evaluated against the antigen (0.01 to 1 μg) using established Western blot analysis procedures^10^. The anti-ABHD14A antibody was purified via affinity chromatography by covalently coupling the antigen as a bait on a SulfoLink resin (buffer used: 50 mM Tris, 5 mM EDTA at pH 8.5). Following coupling and cysteine blocking, the resin was washed extensively, first with 1 M NaCl and subsequently with 20 mM Tris at pH 7.5. The bound anti-ABHD14A polyclonal antibodies were eluted using 100 mM glycine (pH 2.5) in 500 µL fractions, neutralized, pooled, and dialyzed into storage buffer [50% (v/v) glycerol, PBS, 0.2% (w/v) sodium azide]. The antibody performance was validated at a dilution of 1:1000 against various amounts of the antigen (0.01 to 1 μg) relative to a commercially available antibody (Sigma-Aldrich; catalog no: SAB4501087) by Western blotting protocols previously reported by us^10^.

### ABHD14A expression in immortalized cell lines and mouse tissues

The expression of ABHD14A in various mammalian cell lines or adult mouse tissues was assessed using Western blot analysis using establish protocols^10^. Briefly, lysates (50 μg) from several immortalized mammalian cell lines (A549, BV2, HEK293T, HeLa, HepG2, Neuro2A, NIH3T3, RAW264.7, THP-1) or adult mouse tissues (brain, heart, liver, lung, kidney, muscle, pancreas, spleen, testis) were analysed by Western blot analysis using our in house anti-ABHD14A antibody or a commercially available anti-ABHD14A (Sigma-Aldrich; catalog no: SAB4501087) at a dilution of 1:500. In these immunoblotting experiments, 0.05 μg of purified human ΔN60-WT ABHD14A was used as a positive control, and Ponceau staining was used to ensure appropriate protein loading.

### Overexpression of rat ABHD14A variants in HEK293T cells

The full length WT rat ABHD14A (UniProt ID: Q5I0C4) cloned in the pCMV-Sport6 mammalian expression vector was purchased from GE Dharmacon (now Horizon Discovery), and the catalytically inactive S142A variant was generated in the same vector using the site directed mutagenesis strategy (New England Biolabs) as per the manufacturer’s instructions. The rat ABHD14A variants were overexpressed in HEK293T cells using the Lipofectamine 2000 (Thermo Fisher Scientific) via a transient transfection strategy. Briefly, HEK293T cells were cultured in Dulbecco’s Modified Eagle Medium (DMEM) containing 5% (v/v) fetal bovine serum and 1% (v/v) penicillin–streptomycin till 60% confluency. At this confluence, the cells were transfected with DNA (10 µg) in a 1:4 DNA:Lipofectamine ratio in OptiMEM media (Thermo Fisher Scientific), incubated for 45 min prior to addition to cells. Post-transfection, the media was replaced after 12 h, and cells were harvested after 60 h. Lysates were prepared by sonication in PBS, and soluble and membrane fractions were separated by centrifugation (21,000g, 4 °C, 45 min). Gel-based ABPP analysis were performed on the membrane lysates (40 µg) from ABHD14A-transfected or mock-transfected HEK293T cells using FP-rhodamine (10 μM) as per protocols previously reported by us^11^. Overexpression of ABHD14A variants in HEK293T cells was confirmed by Western blot analysis using anti-ABHD14A antibody (rabbit; in house antibody or Sigma-Aldrich; catalog no: SAB4501087; 1:1000 dilution) (primary antibody) and HRP-conjugated anti-rabbit antibody (Goat IgG) (Invitrogen; catalog no: 31460; 1:10000 dilution) (secondary antibody) as per protocols previously reported by us^11^.

### Cellular localization of ABHD14A

The subcellular localization of ABHD14A was assessed in HEK293T cells transfected with full length WT rat ABHD14A, where mock-transfected cells were used as controls. The cells were washed, fixed, permeabilized and blocked as reported earlier^12, 13^, and probed using the following primary antibodies: anti-ABHD14A antibody (rabbit; in house antibody or Sigma-Aldrich; catalog no: SAB4501087; 1:100 dilution), and anti-GM130 antibody (BD Biosciences, catalog no: 610822; 1:100). The nuclei were stained with 4′,6-diamidino-2-phenylindole (2 min, 25 °C) as reported earlier^12, 13^. The secondary antibodies used in this study were: Alexa Fluor-633 anti-rabbit IgG (Invitrogen; catalog no: A21070; 1:1000) and Alexa Fluor-568 anti-mouse IgG (Invitrogen; catalog no: A11004; 1:1000). All slides were imaged on a Zeiss confocal microscope, and analysed using the ImageJ software (version 2.1.0/1.53c)^14, 15^.

## RESULTS

### Purification of ΔN60-ABHD14A

Full length WT human ABHD14A contains 271 amino acids and structurally possesses two domains based on a bioinformatics analysis^9^: a transmembrane helix (amino acids: 32 – 55) and the ABHD region (amino acids: 71 – 268) (**Figure 1**). First, we attempted to express the codon optimized full length WT human ABHD14A recombinantly in *E. coli*, but were unsuccessful, despite screening various *E. coli* competent cells, in combination with changing the position of the 6X-His tag (N- or C-terminus) and temperature for inducing expression. Hence, we decided to sequentially truncate the full length protein by sequentially deleting 10 amino acids at a time from the N-terminal end of human ABHD14A, and recombinantly express these truncated variants in *E. coli*. From this experiment, we found that deleting 50 amino acids from the N-terminal end of human ABHD14A did not have any effect on either expression or solubility of the truncated protein (**Figure 1**). However, removing the entire N-terminal transmembrane helix (deleting 60 or more amino acids from the N-terminus) remarkably improved both the expression and solubility of these truncated variants of human ABHD14A (**Figure 1**). Thus, based on this initial screening, we found that deleting the first 60 amino acids from the N-terminus of human ABHD14A (ΔN60-WT ABHD14A variant) was best suited for subsequent biochemical analysis, as it retained most of the full-length protein sequence, after removal of the N-terminal helix.

**Figure 1.**
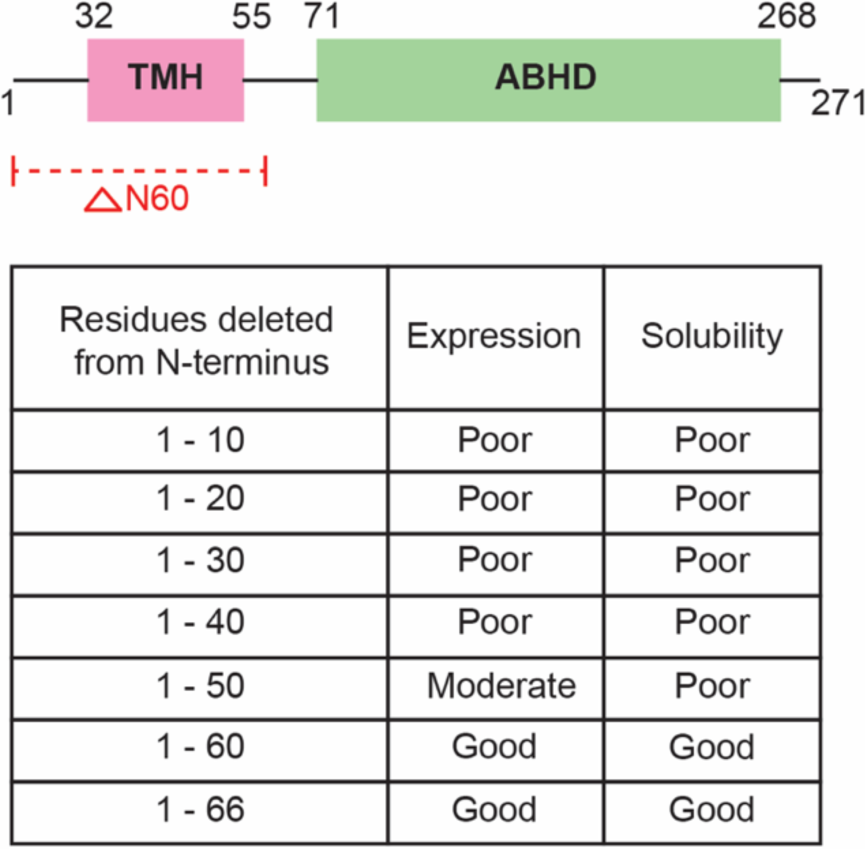
Schematic representation of the human ABHD14A structure. Based on its protein sequence, human ABHD14A is predicted to contain a N-terminal transmembrane helix (TMH) (residues 32 – 55) and an overall α/β-hydrolase domain (ABHD) (residues 71 – 268). The table at the bottom shows the expression and solubility status of various N-terminal truncated variants of human ABHD14A recombinantly expressed in *E. coli*. Amongst the truncated variants, ΔN60-ABHD14A was taken forward for biochemical analysis.

Having confirmed the expression and solubility of the ΔN60-WT ABHD14A variant, next, we purified this truncated protein using affinity chromatography to near homogeneity (> 95% purity) (**Figure S1**), with a typical yield of 5 mg/L culture. The catalytic serine for human ABHD14A is Ser-171, and we generated the ΔN60-S171A ABHD14A variant using site-directed mutagenesis. We purified this mutant to near homogeneity using affinity chromatography (yield 1.5 mg/L culture) and confirmed by analytical gel-filtration that both the truncated ABHD14A variants, ΔN60-WT and ΔN60-S171, behaved alike in this experiment (**Figure S2**).

### Gel-based ABPP assays with ΔN60-ABHD14A variants

To determine activity status of the ΔN60-ABHD14A variants, we first used the established gel-based ABPP assays^10^. Here, we found that as a function of an increasing enzyme concentration (0.2 – 10 μM), ΔN60-WT ABHD14A, but not ΔN60-S171 ABHD14A, displayed robust dose-dependent activity against the FP-rhodamine activity probe (5 μM), which was kept constant in this assay (Figure 2). Along similar lines, we found that as a function of increasing the FP-rhodamine activity probe concentration (0.2 – 5 μM), and keeping the enzyme concentration constant (10 μM), ΔN60-WT ABHD14A again showed robust dose-dependent activity, while the ΔN60-S171 ABHD14A mutant showed no activity at all in this gel-based ABPP experiment (Figure 2). As a control, we used heat-denatured ΔN60-WT ABHD14A (10 μM) in both the gel-based ABPP experiments, and found that unfolding ΔN60-WT ABHD14A resulted in complete loss of enzymatic activity (Figure 2). This established an equivalence between denatured ΔN60-WT ABHD14A and the ΔN60-S171 ABHD14A mutant, and we decided to use denatured ΔN60-WT ABHD14A as a negative control for all subsequent biochemical assays.

**Figure 2.**
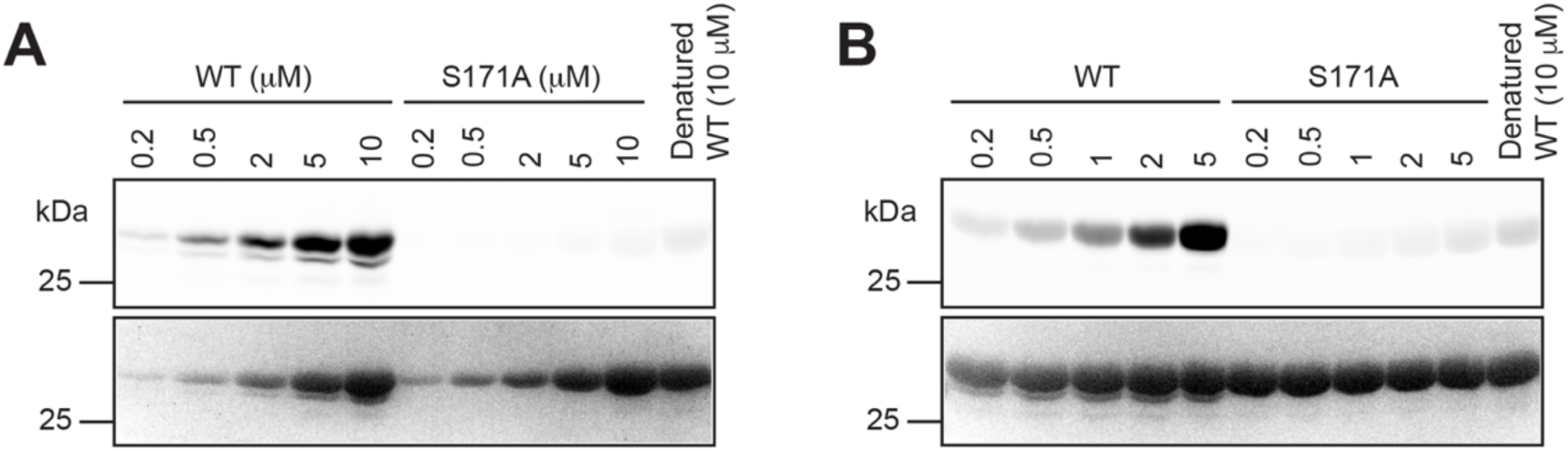
Gel-based ABPP analysis on the ΔN60-ABHD14A variants. (**A, B**) Gel-based ABPP assays on the WT and S171A human ΔN60-ABHD14A variants, showing robust dose-dependent activity of ΔN60-WT ABHD14A, but not ΔN60-S171A ABHD14A, as a function of increasing (**A**) enzyme concentration (0.5 – 10 μM) or, (**B**) activity probe (FP-rhodamine) concentration (0.2 – 5 μM), respectively, while keeping the other constant [FP-rhodamine (5 μM) in panel A, enzyme concentration (10 μM) in panel B]. The gel-based ABPP experiments were performed three independent times with reproducible results each time.

### Acetyltransferase activity of ΔN60-WT ABHD14A

To complement the gel-based ABPP assays, we also performed substrate hydrolysis assays with p-nitrophenyl-esters using established protocols^10^. Here, we incubated native ΔN60-WT ABHD14A and its denatured form with p-nitrophenyl-acetate (pNP-acetate) (500 μM). From this assay, we found that native ΔN60-WT ABHD14A, but not its denatured form, efficiently produced *p*-nitrophenolate from pNP-acetate (approximately 5-fold better) (Figure 3). We also tested other long acyl-chain containing p-nitrophenyl-esters (e.g. butyrate, octonoate, decanoate, dodecanoate, palmitate), and found that ΔN60-WT ABHD14A was unable to produce *p*-nitrophenolate from any of these under similar assay conditions.

**Figure 3.**
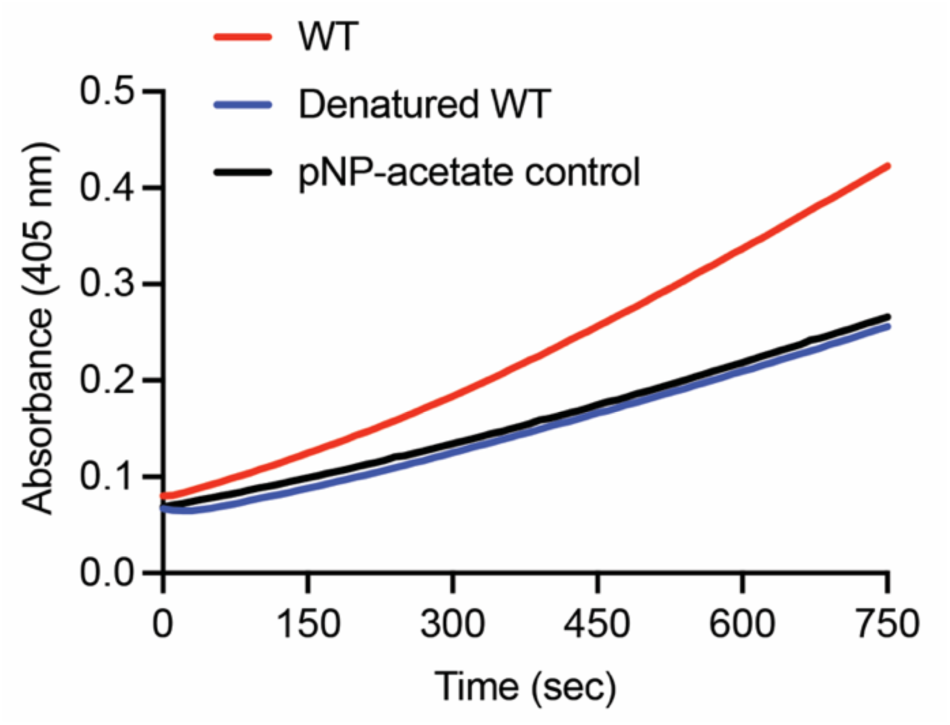
ΔN60-WT ABHD14A can hydrolyze pNP-acetate. Colorimetric enzymatic assay showing ∼5-fold more activity of the native ΔN60-WT ABHD14A, compared to its denatured form, against pNP-acetate (500 μM). The colorimetric assays were performed three independent times with reproducible results each time.

We have previously shown that a closely related mammalian protein ABHD14B possesses a unique ping-pong type acetyltransferase activity with Co-A being the acetyl-group acceptor, and functions as a lysine deacetylase in mammalian cells and tissues^10, 16^. Given the high sequence and structural homology between the two ABHD14 proteins^9^, we wanted to check if ΔN60-WT ABHD14A also possesses any acetyltransferase activity. To test this premise, ΔN60-WT ABHD14A (10 μM) was incubated with Co-A or acetyl-Co-A (1 mM each), and the reaction was initiated by adding pNP-acetate (50 μM). Consistent with an acetyltransferase activity, we found that Co-A, but not acetyl-Co-A, significantly increased the rate of pNP-acetate hydrolysis (Figure 4). Consistent with findings from experiments with human ABHD14B^10, 16^, this activity profile is evidence for the formation of acetyl-Co-A from pNP-acetate with Co-A being the acetyl-group acceptor (heightened acetyl-Co-A formation in the experiment was also verified by LC-MS analysis). Overall, our studies show that ΔN60-WT ABHD14A indeed possesses an acetyltransferase activity similar to its close mammalian homolog ABHD14B.

**Figure 4.**
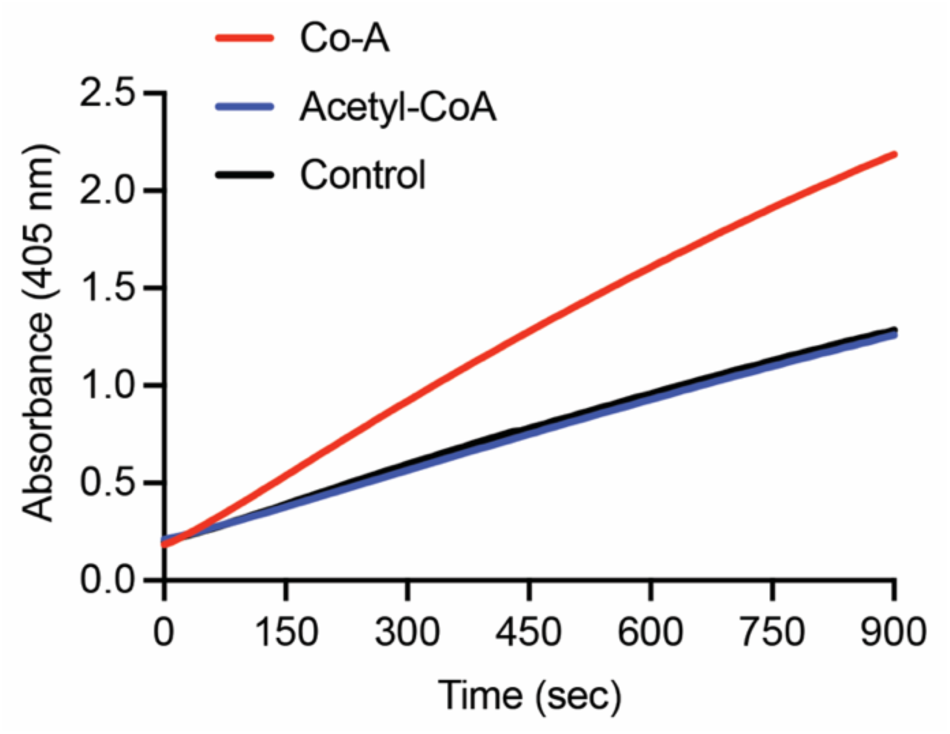
ΔN60-WT ABHD14A has acetyltransferase activity. Colorimetric enzymatic assay with pNP-acetate (50 μM), showing an increase in the rate of enzymatic reaction following incubation of ΔN60-WT ABHD14A with Co-A, but not acetyl-Co-A. This is consistent with an acetyltransferase reaction that results in the formation of acetyl-Co-A, when Co-A and pNP-acetate are incubated with ΔN60-WT ABHD14A. The colorimetric assays were performed two independent times with reproducible results each time.

### Characterization of an anti-ABHD14A antibody

Since we were successful in purifying ΔN60-WT ABHD14A in good yield and high purity, we decided to generate an anti-ABHD14A antibody (polyclonal) from a single rabbit using in house protocols at the NFGFHD at IISER Pune. Post-immunization and the terminal bleed, we obtained 30 mL of serum, a fraction of which (12 mL of serum) was used to purify approximately 3.5 mg of anti-ABHD14A polyclonal antibody from one rabbit. To validate the compatibility in Western blotting experiments, we first tested the serum (from the terminal bleed) and purified antibody against varying amounts (0.01–1 μg) of ΔN60-WT ABHD14A and ΔN60-S171A ABHD14A. From this Western blot analysis, we found that both the serum and purified polyclonal antibody at a dilution 1:1000 could reliably detect 50 ng of ΔN60-ABHD14A variants (Figure 5), thus, giving us confidence that this in-house generated polyclonal antibody might in fact be useful in detecting endogenous ABHD14A (if present) in immortalized mammalian cell lines and adult mouse tissues by Western blotting.

**Figure 5.**
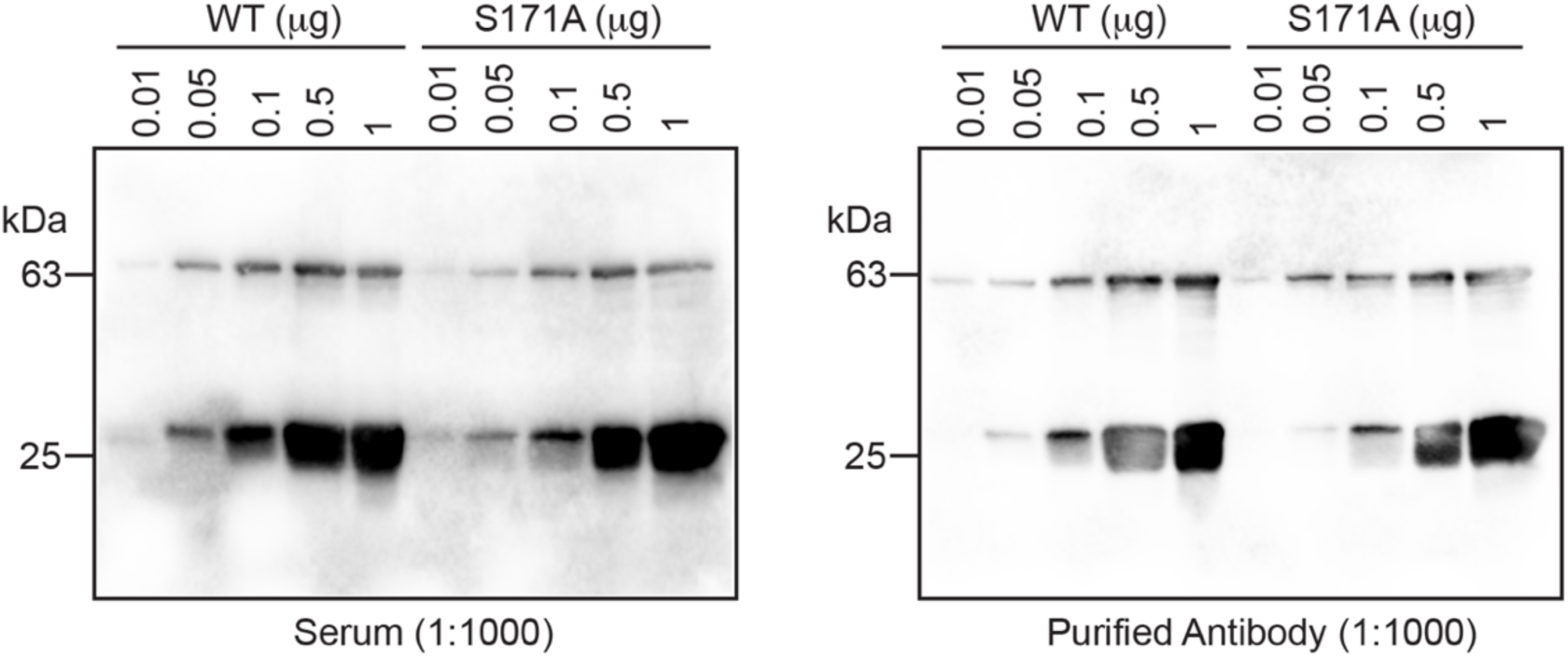
Characterization of the anti-ABHD14A antibody. Western blot analysis of a rabbit polyclonal anti-ABHD14A antibody (*left panel*, serum after terminal bleed at a dilution of 1:1000; *right panel*, purified antibody at a dilution of 1:1000) tested against varying amounts (0.01 – 1 μg) of recombinantly purified ΔN60-WT ABHD14A and ΔN60-S171A ABHD14A. All Western blot experiments were performed three independent times with reproducible results each time.

Several large-scale gene expression databases suggest that ABHD14A is present in various mammalian cell lines and adult mouse tissues^17–21^. However, in comparison, there are very limited studies that quantify the ABHD14A protein levels in any mammalian cell lines or tissues. To investigate this, we tested our anti-ABHD14A polyclonal antibody and surprisingly found that ABHD14A was absent in all the immortalized mammalian cell lines and adult mouse tissues tested by us (Figure 6). To confirm that this result wasn’t an artefact of our antibody, we also purchased a validated commercially available antibody and found similar results in the same Western blot analysis (Figure 6). To ensure that all the Western blot experiments were working well, we also used 50 ng of ΔN60-WT ABHD14A as a positive control in these immunoblotting experiments and were able to detect very good signals for this positive control in all the experiments (Figure 6).

**Figure 6.**
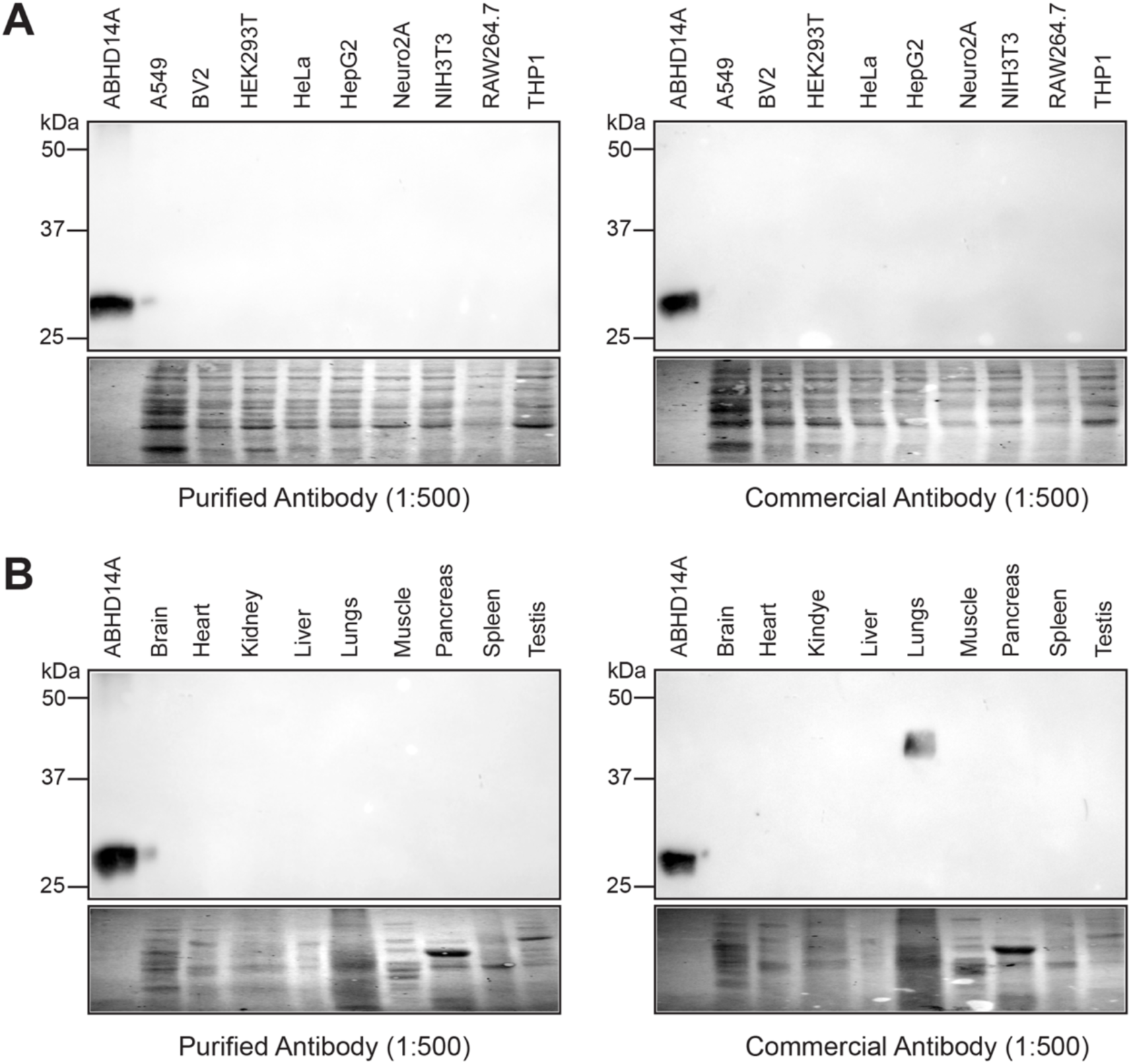
Detection of ABHD14A in mammalian cells and mouse tissues by immunoblotting. (**A, B**) Western blot analysis with our rabbit polyclonal anti-ABHD14A antibody and a commercially available rabbit polyclonal anti-ABHD14A antibody tested against (**A**) proteomes (50 μg) of different immortalized mammalian cell lines, and (**B**) proteomes (50 μg) of different adult mouse tissues. The Western blots show that ABHD14A is not expressed (or below detection levels) in any of the immortalized cell lines or adult mouse tissues tested. The ΔN60-WT ABHD14A (50 ng) was used as a positive control in all the immunoblotting experiments. All Western blot experiments were performed three independent times with reproducible results each time.

Taken together, this result suggests that contrary to information available from large-scale gene expression databases, ABHD14A is perhaps physiologically produced on demand or in response to specific stimuli and/or environmental conditions in mammalian cells and tissues.

### Expression and localization of ABHD14A in HEK293T cells

Since we were unable to find a suitable immortalized mammalian cell line to study endogenous ABHD14A, we decided to (over)express and map the subcellular localization of rat ABHD14A in the mammalian HEK293T cell line. The human and rat ABHD14A proteins have very high sequence similarity (87%) and complete conservation of residues from the N-terminal transmembrane helix and the ABHD region of the polypeptide sequence^9^. Hence, both proteins are likely to behave the same in these experiments and therefore, we decided to use the rat ABHD14A for the experiments in HEK293T cells. Using a lipofectamine-based transient transfection strategy, we expressed WT and the catalytically inactive S142A variant of rat ABHD14A in HEK293T cells. Relative to a mock control, the overexpression of both WT and S142A variant of rat ABHD14A in the membrane lysates of HEK293T cells was confirmed by Western blot analysis using our anti-ABHD14A polyclonal antibody (Figure 7). Consistent with the biochemical experiments described earlier with the ΔN60-ABHD14A variants, we found that relative to the mock control, WT rat ABHD14A, but not the corresponding catalytically inactive S142 mutant, showed robust activity in the membrane lysates of HEK293T cells by gel-based ABPP analysis (Figure 7). This result conclusively shows that full-length mammalian ABHD14A is indeed an active enzyme, when (over)expressed in mammalian cells.

**Figure 7.**
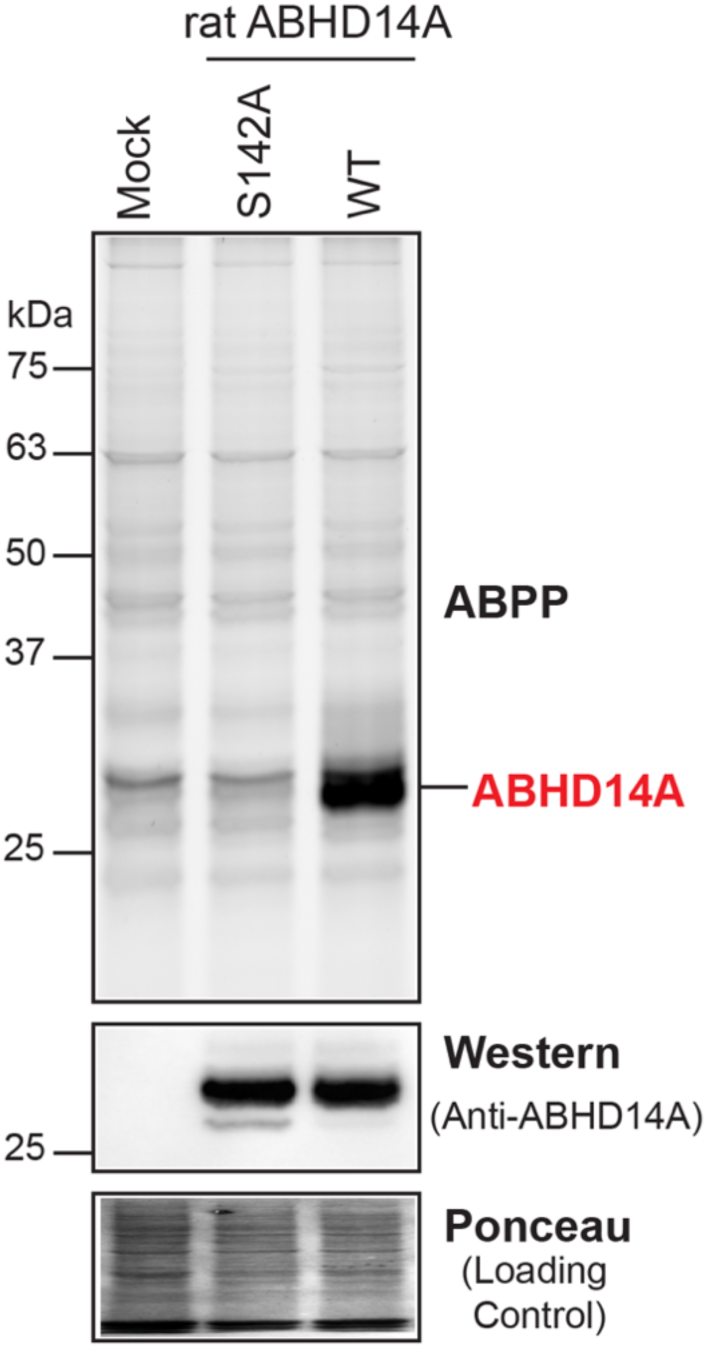
Activity of full length rat ABHD14A in HEK293T cells. Membrane proteomes (50 μg) of HEK293T cells transfected with mock, WT rat ABHD14A, or S142A rat ABHD14A were assessed by gel-based ABPP (top panel), Western blot analysis with our anti-ABHD14A antibody (middle panel), and Ponceau S staining (bottom panel). This experiment was done 3 times with reproducible results each time.

Full length mammalian ABHD14A possesses a conserved N-terminal transmembrane helix that presumably anchors this enzyme to the membrane of some cellular organelle. To assess the subcellular localization of ABHD14A, we performed an immunofluorescence analysis (IFA) on HEK293T cells that (over)expressed WT rat ABHD14A using our anti-ABHD14A polyclonal antibody. We found from this cellular IFA, that cellular fluorescent signal for ABHD14A (red channel) was only visible when ABHD14A was (over)expressed in HEK293T cells, and the mock control samples showed negligible signal for ABHD14A (Figure 8). This experiment validated that our anti-ABHD14A antibody was also compatible for cellular IFA experiments. Quite interestingly, (over)expressed ABHD14A showed a dense aggregation pattern, consistent with proteins that localize to the Golgi apparatus. Indeed, we found that the intense cellular fluorescence signal for ABHD14A (red channel) highly co-localized with the cellular fluorescence signal for a bonafide Golgi apparatus marker, GM130 (green channel) (Figure 8). We also performed cellular immunofluorescence experiments with other organellar markers (e.g. endoplasmic reticulum, plasma membrane, mitochondria), but did not find any co-localization with ABHD14A, suggesting that ABHD14A is predominantly localized to the Golgi apparatus, when expressed in mammalian cells.

**Figure 8.**
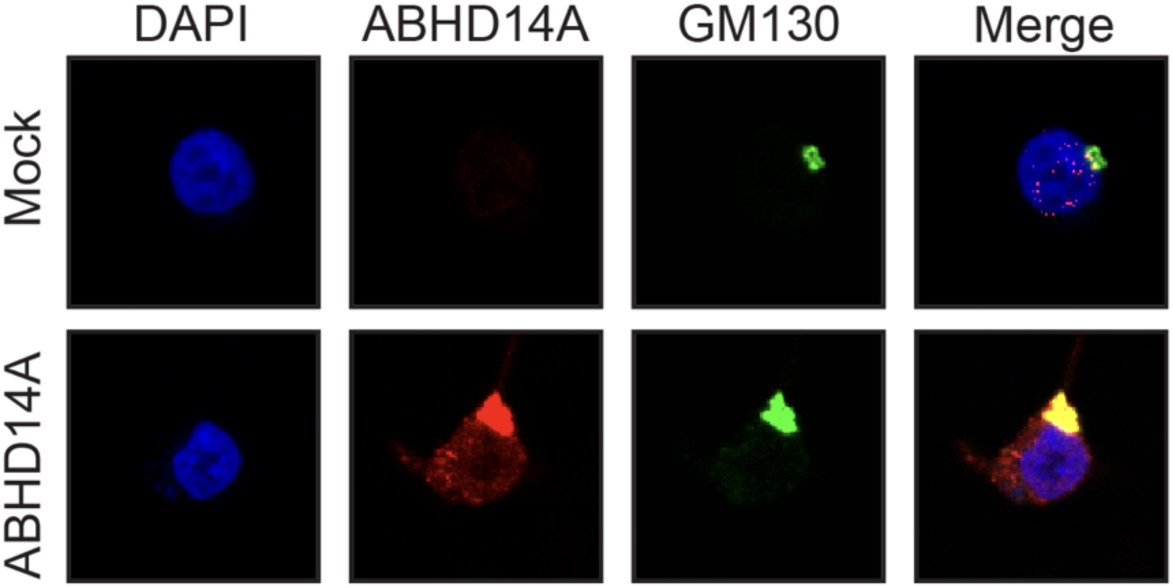
Cellular localization of (over)expressed ABHD14A in HEK293T cells. A cellular immunofluorescence analysis (IFA) in HEK293T cells shows that (over)expressed ABHD14A is present in Golgi-apparatus, and the cellular fluorescence for ABHD14A (in the red channel) is seen only when ABHD14A is (over)expressed in HEK293T cells. The cellular IFAs were performed three independent times with reproducible results each time.

## DISCUSSION

This study provides the first functional evidence that ABHD14A is an active mSH with acetyltransferase capacity, and it brings clarity to a long-standing gap in the annotation of this enigmatic enzyme. Until now, ABHD14A had been known almost exclusively from sequence predictions and structural modelling^9^, with no consensus on its catalytic activity, substrate range, or cellular localization. By integrating biochemical assays with newly developed immunological tools and a mammalian expression system, our findings enable ABHD14A to be placed more concretely within the mSH family, while simultaneously revealing surprising features that distinguish it from established members of this enzyme class.

One conceptual advance emerging from this work is the clear functional similarity between the two ABHD14 proteins, namely ABHD14A and ABHD14B^9^. Although the two proteins share only moderate sequence identity^9^, their catalytic behaviour appears mechanistically aligned, particularly the enhancement of short-chain ester turnover in the presence of Co-A. This pattern is characteristic of ping-pong type acetyltransferase chemistry^10, 16^, suggesting that ABHD14A may also participate in Co-A linked acetyl-group exchange reactions in mammalian cells. What is interesting is that ABHD14A and ABHD14B have distinct subcellular localizations (Golgi apparatus for ABHD14A, and Cytosol and nucleus for ABHD14B^10^), and perhaps, these enzymes access different yet distinct substrates. Yet, in both cases, the restricted activity towards pNP-acetate, implies a finely tuned active site that favors small acetyl donors rather than broader lipid hydrolysis. This places ABHD14A (along with ABHD14B) in a unique niche within the mSH family^5^: an enzyme structurally classified among hydrolases, yet operating with functional elements of a transferase. Such mechanistic hybridity has been observed for only a handful of human mSHs, underscoring its biological rarity and importance.

A second major insight derives from our protein-level expression analyses. While public transcriptomic and proteomic resources consistently report widespread ABHD14A expression in human tissues and cells^17–21^, our experimental data challenge these predictions. Using two independent antibodies, including a rigorously validated polyclonal reagent generated in this study, we were unable to detect endogenous ABHD14A in any immortalized cell line or adult mouse tissue examined. This discrepancy highlights limitations of high-throughput resources, where detection thresholds, peptide inference issues, or reliance on mRNA abundance can lead to overestimated protein prevalence^22, 23^. The absence of protein-level detection suggests that ABHD14A expression is either extremely low under basal conditions or is tightly regulated and induced only in specific physiological contexts. Such context-dependent expression has been documented for several mSHs involved in stress responses, lipid remodeling, and immune activation, raising the possibility that ABHD14A plays a similarly specialized role that is not captured in standard culture or steady-state tissue conditions. Our cellular experiments provide additional clues about potential physiological functions.

When expressed in HEK293T cells, full length ABHD14A localizes predominantly to the Golgi apparatus, distinguishing it from many mSHs that reside in the cytosol or endoplasmic reticulum. Golgi-localized hydrolases or transferases are often involved in lipid maturation, vesicular trafficking, glycoprotein processing, or regulation of secretory pathways^24, 25^. The dense punctate signal observed for ABHD14A is characteristic of enzymes associated with Golgi membranes and suggests that its functional substrates may include proteins or metabolites trafficking through this secretory system^26, 27^. Whether ABHD14A acts upstream (e.g., regulating acetyl-CoA pools at the Golgi), downstream (e.g., modifying lumen-facing proteins), or in parallel with canonical Golgi enzymes remains an open question. Importantly, the catalytic inactivity of the S142A mutant in both recombinant assays and cellular ABPP confirms that the endogenous activity observed in cells is intrinsic to ABHD14A and not attributable to off-target effects associated with its overexpression.

Beyond the biochemical and cellular findings, this study also provides valuable methodological tools that will enable future work. The successful expression of a soluble N-terminally truncated variant overcomes a long-standing barrier to its biochemical characterization. The newly generated high-affinity anti-ABHD14A polyclonal antibody now makes it possible to screen physiological conditions, developmental stages, or disease states in which ABHD14A expression may be induced. Together, these tools set the stage for more targeted investigations that extend well beyond what was previously feasible.

## FUTURE DIRECTIONS

Several important directions arise from this work as we move toward determining the physiological relevance of ABHD14A. A key priority is the identification of endogenous substrates or interacting proteins, particularly in light of the enzyme’s acetyltransferase activity and Golgi localization. Chemical proteomics approaches such as ABPP, CoA-reactive probe profiling, or substrate-trapping mutants will be essential for mapping the molecular environment in which ABHD14A operates. Equally important will be establishing the physiological contexts in which ABHD14A expression is induced. A previously reported study shows a transcriptomic link between ABHD14A and the transcription factor Zic1, a well-characterized regulator of early brain development^28^. The co-expression patterns of these genes in neurogenic niches coupled to population wide gene association analysis suggest that ABHD14A may participate in developmental programs that shape neuronal identity, maturation, or Golgi-dependent processing of neurodevelopmental signalling molecules, and their associations with neurological diseases^28–32^. To explore these possibilities, model systems like neural progenitor differentiation assays, mouse embryonic brain tissue, or human brain organoids could be leveraged to profile ABHD14A expression dynamics relative to Zic1 activity. It will be particularly informative to determine whether ABHD14A influences Zic1-regulated transcriptional landscapes indirectly, perhaps by modulating acetyl-CoA availability, Golgi-based processing of secreted cues, or post-translational modifications of proteins involved in neurodevelopmental signalling. High-resolution structural studies and genetic manipulation strategies, including knockout or knock-in models, will further clarify whether ABHD14A serves as a biochemical effector downstream of Zic1-regulated developmental pathways. Collectively, these future efforts promise to uncover whether ABHD14A represents a previously unrecognized enzymatic node linking mSH chemistry to the molecular architecture of brain development.

## CONCLUSIONS

This study provides the first clear biochemical and cell-based evidence that ABHD14A is an active member of the mSH family, possessing an acetyltransferase activity. By overcoming long-standing challenges in recombinant expression, we identify a soluble, catalytically competent truncated variant that enabled mechanistic studies, including the discovery of a CoA-dependent acetyltransferase activity. The generation of a robust polyclonal antibody further allowed us to probe endogenous expression, revealing that ABHD14A is absent from commonly used immortalized mammalian cell lines and adult mouse tissues despite predictions of broad transcriptional presence. Its selective Golgi localization upon heterologous expression suggests a specialized physiological role within secretory or membrane-associated pathways. Collectively, these findings not only establish ABHD14A as an active and biochemically distinct enzyme, but also provide foundational tools and mechanistic insights that will guide future efforts to map its endogenous substrates, regulatory cues, and physiological relevance.

## ACCESSION CODES

The UniProt ID of human and rat ABHD14A are Q9BUJ0 and Q5I0C4 respectively.

## AUTHOR CONTRIBUTIONS

S.G. performed all the experiments. S.G. and S.S.K. conceived the project, analyzed the data and wrote the paper. S.S.K. supervised and acquired funding for the project.

## DATA AVAILABILITY

All data the supports the findings of this study are available in the paper and its supporting information or are available with the corresponding author upon reasonable request.

## ACKNOWLEDGEMENTS

This study was financially supported by an EMBO Young Investigator’s Award (to S.S.K.), and a Prime Minister’s Research Fellowship (graduate student fellowship to S.G.). Dr. Swapnil Bangar from the NFGFHD, IISER Pune is thanked for the help with the antibody production. Members of the S.S.K. lab are thanked for providing critical inputs throughout the course of this study.

