## Supplementary Figures for "Biochemical characterization of ABHD14A, an outlying member of the metabolic serine hydrolase family"

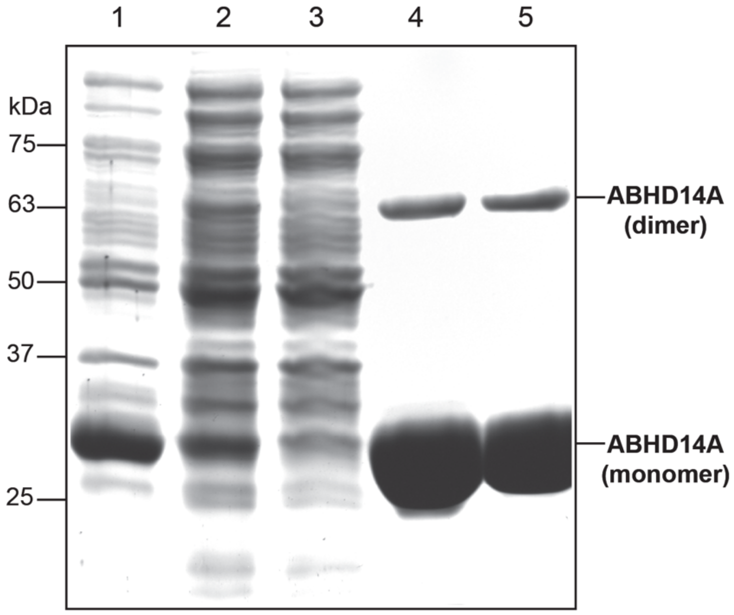


**Figure S1. Purification of ∆N60-WT ABHD14A.** A representative coomassie gel for the scheme towards purifying ∆N60-WT ABHD14A recombinantly from *E. coli*. The monomer and dimer bands of ∆N60-WT ABHD14A were verified for identity using in-gel proteomics analysis. 1 = Whole cell lysate; 2 = supernatant after sonication; 3 = flow through after Ni-NTA column; 4 = elute from Ni-NTA column; 5 = final protein after dialysis.

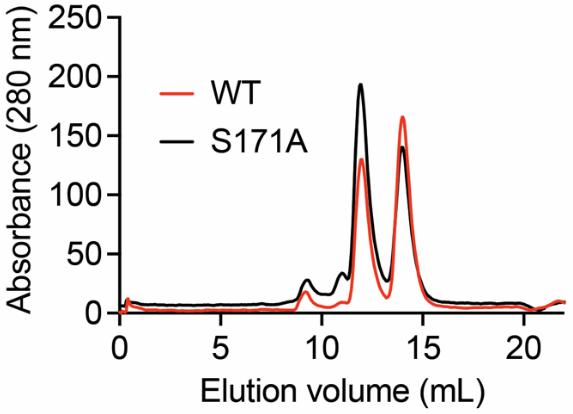


**Figure S2. Analytical gel filtration of ∆N60-ABHD14A variants.** Representative UV-traces from an analytical gel filtration experiment showing that ∆N60-WT ABHD14A and ∆N60-S171A ABHD14A have similar oligomerization states and in turn, tertiary structures. These assays were done three independent times with reproducible results each time.
